# Prediction and discovery of protein-protein direct interactions and stable complexes based on gene co-expression and co-evolution

**DOI:** 10.1101/2024.10.05.616780

**Authors:** P. Miglionico, Lorenzo Amir Nemati Fard, C.J. Gloeckner, F. Raimondi

**Author notes:** **Corresponding authors:** Francesco Raimondi.

## Abstract

In this study we employed a data-driven approach to explore the evolutionary and genetic determinants of protein direct interactions and stable complex formation in the human proteome. We found that simple co-evolutionary and co-expression metrics are highly informative of direct interactions and stable complexes. We used this information to train supervised binary classifiers to predict interactions either directly involved in the formation of a complex (as annotated in IntAct) or forming stable complexes (from Complex Portal). In the former task, our model was able to discriminate direct interactions with an AUROC=0.813, while in the latter it discriminated interaction forming stable complexes with an AUROC=0.964. In both cases, our approach outperformed String, that we employed as a baseline. Feature importance analysis revealed different contributions to the prediction of these distinct interaction types. Co-evolutionary features, in particular those referred to protein domains involved in interaction interfaces, are more important to discriminate direct interactions. On the other hand, co-expression features contributed more to the prediction of stable complexes. From these pairwise predictions we generated a proteome-wide network that we clustered to assess the recovery of known complexes from Complex Portal within network communities. We were able to recover known complexes at a higher accuracy compared to other approaches.

In conclusion, we propose a new method able to discriminate direct interactions as well as forming stable complexes. This method can be used to stratify molecular interaction networks, as well as to perform discovery of new functional complexes at a proteome-wide scale.

## Introduction

Protein-protein interaction networks orchestrate the structure and functioning of the cell and are often disarranged in disease. Advancement in mass spectrometry techniques coupled with high-throughput screenings such as yeast two-hybrid (Y2H) (Luck et al., 2020), affinity-purification (AP-MS) (Huttlin et al., 2021), or proximity labeling (Sears et al., 2019) enabled the discovery of interactions on a proteome-wide scale. However, these experimental approaches produce large interactomic datasets that are often affected by non-specific interactions which do not reflect physiological conditions, leading to high false positive rates. Available resources such as IntAct (del Toro et al., 2022) are engaged in classifying interactions into different types, i.e. direct, physical or association, via a curation process which entails the integration with other experimental sources, including analysis of 3D complex structures. However, the discrimination of direct interactions from indirect or spurious associations within large PPI networks remains a challenging task, often preventing a deeper mechanistic interpretation of these datasets.

Computational strategies to structurally dissect molecular interaction networks have been recently proposed, sparked by the success in AI-driven structural predictions. For instance, AlphaFold-multimer(Evans et al., 2022) has been used to predict with high confidence the structures of thousands of human protein interactions (Burke et al., 2023). Variations of the RosettaFold method (i.e. Rosetta 2 Track (Humphreys et al., n.d.) and RoseTTAFold2-Lite (Humphreys et al., 2024)) have been employed as strategies to screen large PPI network sets to structurally predict, in combination with AF-multimer, the structures of high confidence complexes. Machine learning methods have also been developed to discriminate direct interactions by taking AF-multimer predictions as input (Schmid & Walter, n.d.).

These approaches rely in first place on the structural predictions of the complexes through AI models such as AF-multimer, which are computationally intensive and might be prohibitive at the proteome scale. For this reason, such methods mostly rely on modeling binary interactions, not considering the modeling of higher order multimers. Moreover, given the stringency of the metrics employed, which are based on AF-multimer’s confidence score, these approaches tend to focus only on a limited number of higher confidence structures, opening the possibility of high false negatives rates.

Hence, an algorithm able to process an input interactomic dataset, to score interactions that are more likely to be direct or to for stable complexes, would be highly desirable and could be used to streamline effort to structurally model proteome-wide interactomes.

## Results

### Gene co-evolution and co-expression inform about protein direct interactions and stable complexes

We derived a set of pairwise gene co-evolutionary and co-expression features and checked for their statistical associations with molecular interaction types (Figure 1).

**Figure 1.**
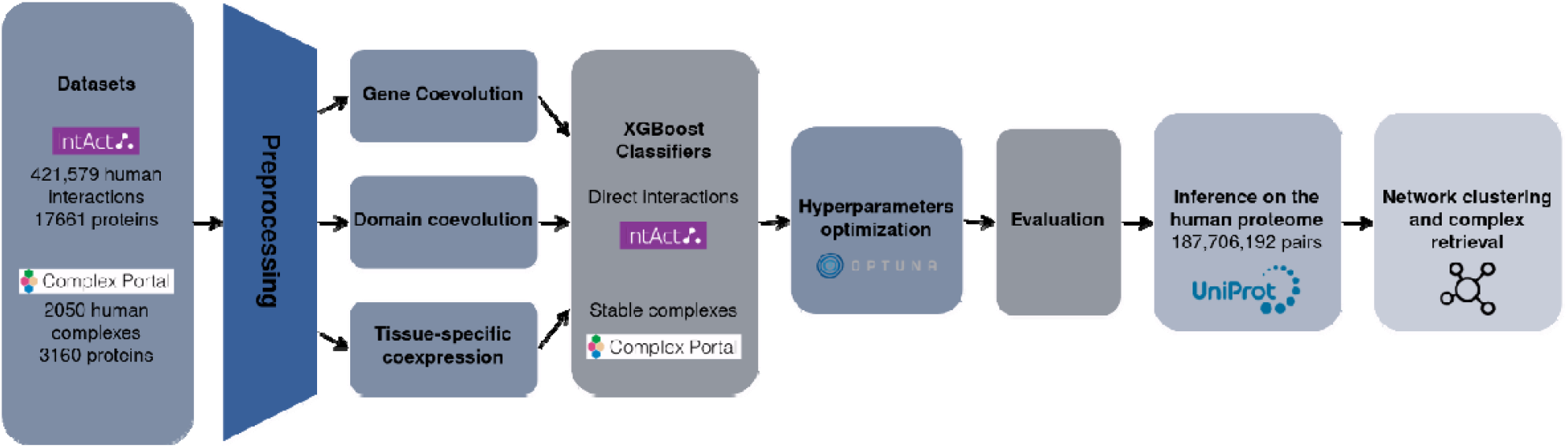
workflow of the procedure. Schematic workflow of the pipeline from dataset acquisition and model training through evaluation and human proteome level clustering. We used the OMA database to identify ortholog sequences, InterPro to identify domains and GTEX and TCGA to compute the Tissue-specific coexpressions. Two model types are trained: one to predict if a protein pair is part of a stable protein complex and one to discriminate direct interactions from weaker associations. After hyperparameter optimization and evaluation, the models are applied to the entire human proteome. The resulting interaction scores are used to construct a weighted network, which is then clustered to reveal protein complexes.

To obtain co-evolutionary features, we first retrieved the list of orthologs for each human gene available from a reference orthology database (i.e. OMA (Altenhoff et al., 2021); see Methods). We estimated the co-evolution between two genes as the degree of their co-presence in sequenced genomes, which we assessed through both Jaccard and Mutual Information metrics (see Methods). We also derived similar features at the domain level by considering the coevolution of pairs of annotated domains (i.e. Interpro (Blum et al., 2021)) on protein pairs across sequenced genomes (see Methods). We considered every possible domain pair as well as those found in spatial contact in 3D structures from the Protein Data Bank (PDB) (see Methods).

Next, we assessed the degree of co-expression of interacting genes from 47 healthy (GTEx) as well as from 32 cancer (TCGA) tissues, using weighted gene co-expression network analysis (WGCNA, (Langfelder & Horvath, 2008)) from which we considered both expression correlation and Topological Overlap Measure (TOM) as an estimate of gene pairs coexpression (see Methods).

We checked for correlation among features and, as expected, we found out that features of the same type (i.e. co-evolutionary or co-expression) have higher correlation values (Figure 2A). Among co-expression features, we found that correlations in healthy sub-tissues preserved the tissue of origin (Figure 2A). Some of them, such as brain or gastro-intestinal tract healthy tissues, are characterized by higher correlation values (Figure 2A). On the other hand, the correlation values derived from cancer tissues were characterized by higher values, irrespective of the tissue of origins, which suggests that tumor tissues have lost their original transcriptional program characteristic of the healthy tissue to foster oncogenic transformation.

**Figure 2.**
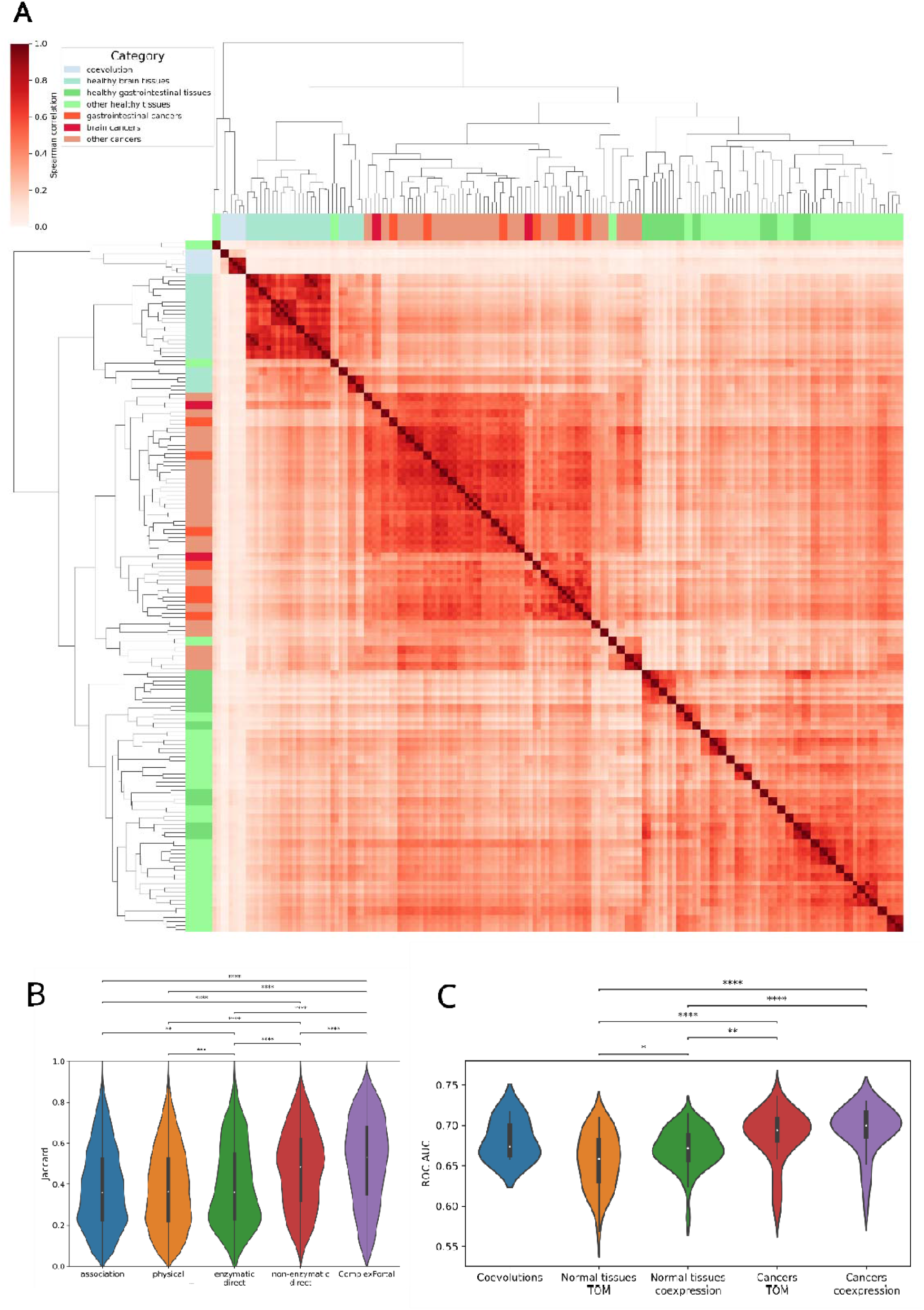
Analysis of Individual Features A) Clustermap of the pairwise Spearman correlations of the 164 features; B) Distribution of the Jaccard coevolution across all the PPIs annotated in Intact; C) Area under the ROC curve of the different feature categories in normal and tumor tissues for the task of identifying protein pairs involved in stable complexes. P-values refer to a Mann-Whitney test with Bonferroni correction (*p<0.05, **p<0.01, ***p<0.001, ****p<0.0001).

We then used the co-evolution and co-expression values as features (independent variables) of gene pairs, and we employed the interaction types as target variables (i.e. classification labels). We considered two protein-protein interaction (PPI) datasets. First, we used the entire human interactome from IntAct ((del Toro et al., 2022), i.e. IA set), where we stratified interactions according to the annotated interaction type, i.e. Association, Physical and Direct (in turn sub-divided into Enzymatic and Non-enzymatic) interactions. To these standard annotated types, we added an additional term depending on whether the interaction is reported to participate in stable complexes from Complex Portal (Meldal et al., 2022). We also generated a second dataset by considering all the human interactions from Complex Portal, which we combined with randomly picked protein-protein pairs which we considered as a negative, background set (i.e. CP set; see Methods).

We found that distinct interaction classes from IntAct are associated with significantly different distributions of these features (Figure 2B). For instance, features based on coevolution have significantly higher values for stable complexes or direct interactions (Figure 2B) compared to other interaction types (i.e. Association or Physical interaction). In particular, all 164 features (both coevolution and coexpression based) were significantly higher in interactions between proteins belonging to the same complex (from Complex Portal, with maximum P-value < 10^-58, Mann-Whitney test with Bonferroni correction on the IA set).

We found that certain individual features were predictive of certain interaction types. In the IA set, we found 42 unique features with AUROC values greater than 0.7 for interactions forming stable complexes in Complex Portal, with coexpression from lung adenocarcinoma (“*lung_adenocarcinoma_TOM”)* being the single feature with the highest AUROC (0.736) (Supplementary Table 1). In general, we found higher AUROC values for co-expression features over the co-evolutionary ones for the prediction of stable complex interactions. In particular, co-expressions from cancer transcriptomics are characterized by higher AUROC for the prediction of stable complexes (Figure 2C), suggesting a higher activity of complexes sustaining oncogenic processes.

### Training a stable interaction classifier based on co-evolutionary and co-expression features

We employed 6 coevolutionary and 158 coexpression features to train a supervised machine learning algorithm to discriminate interactions involved in stable complexes and direct interactions. We employed a state of the art algorithm for supervised learning on tabular feature sets (i.e. XGBoost (Chen & Guestrin, 2016), with a bayesian procedure for optimal hyperparameters search (i.e. Optuna (Akiba et al., 2019)).

The model is able to discriminate interactions forming stable complexes in the CP dataset with AUROC 0.964 (Figure 3A). In the IA dataset, the model achieves an AUROC of 0.920 for the same task. The slightly worse performance on the IA dataset is likely attributed to using protein pairs froma other interaction types (i.e. Physical and Associations) as negatives instead of random pairs. To further assess the model’s capacity to identify novel complexes, we randomly selected 10% of the complexes in Complex Portal as a “held-out” set. Proteins within these complexes were excluded from the training data, and the model was trained on the remaining CP set. We then evaluated the model’s ability to retrieve stable interactions within this held-out protein set, achieving an AUROC of 0.802 on this task.

**Figure 3.**
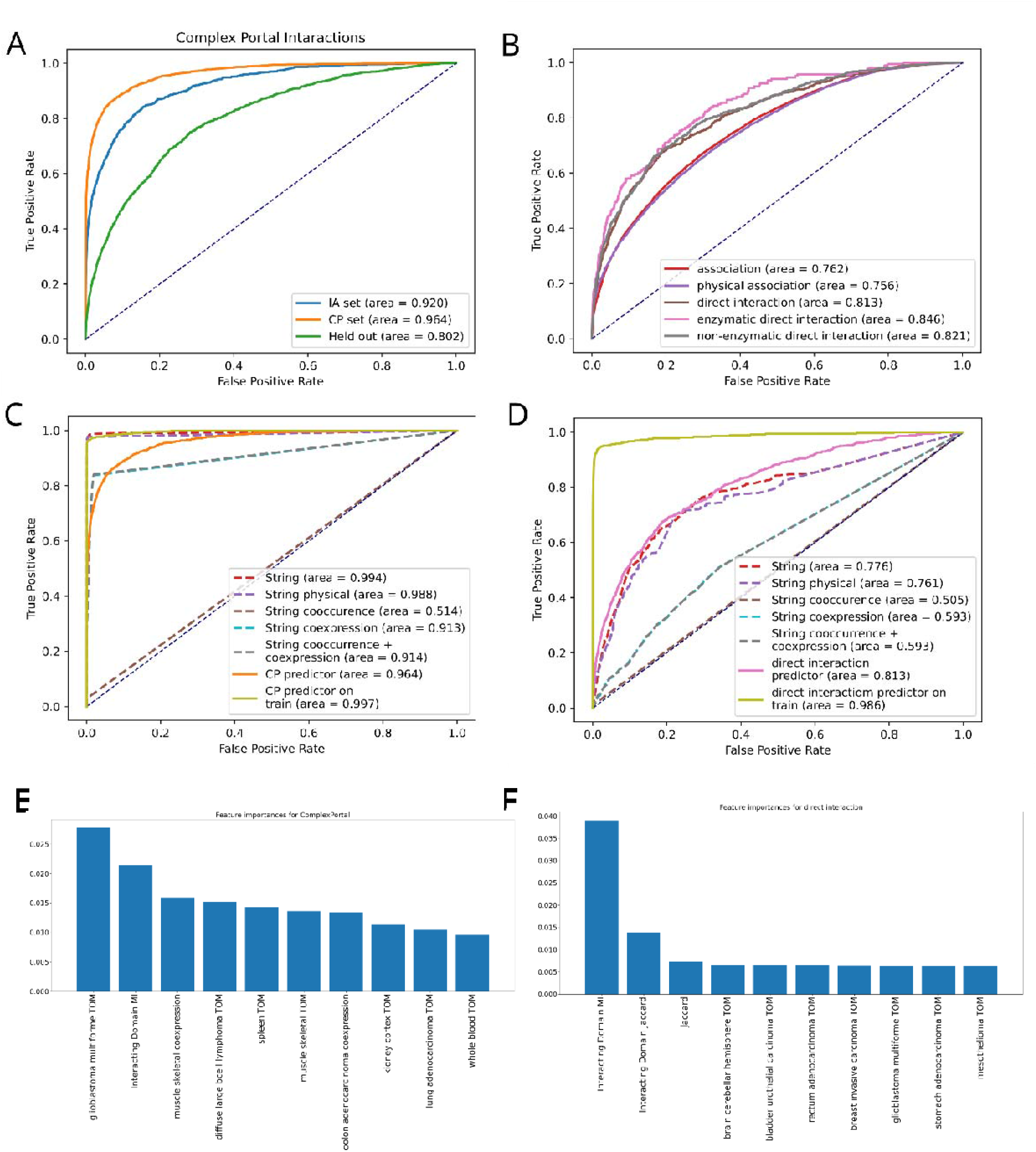
Machine learning models to predict direct and stable complex interactions. A) ROC curves of the model trained to discriminate interactions forming stable complexes on the IA set (blue), on the CP set (orange), on a set of random complexes excluded from the training set in their entirety (green); B) ROC curves of the models trained on the IA set to discriminate associations (red), physical associations (purple), direct interactions (brown), enzymatic direct interactions (pink), non-enzymatic direct interactions (gray); C) ROC curves of the model trained to discriminate interactions forming stable complexes on the CP set (orange) compared to the ones obtained using the different STRING confidence scores as predictors for the same task; D) ROC curves of the model trained to discriminate direct interactions on the IA set (pink) compared to the ones obtained using the different STRING confidence scores as predictors for the same task; E) Histogram representing the 10 most important features for the task of discriminating interactions forming stable complexes; F) Histogram representing the 10 most important features for the task of discriminating direct interactions. Feature importances are computed as the average loss change due to each feature.

We additionally utilized the IA set to train a model for predicting the five identified interaction types: association, physical association, non-enzymatic direct interaction, and enzymatic direct interaction, direct interaction (the union of non-enzymatic and enzymatic direct interactions). The ROC curves for association and physical association predictors are approximately 0.76, while those for the three direct interaction categories range between 0.81 and 0.85 (Figure 3B).

We compared the performance of trained models with one of String networks on the entire human Complex Portal as well as IntAct datasets. On the CP dataset, our best performing model performed similarly to STRING and STRING Physical (Szklarczyk et al., 2023). It must be noted, however, that while String considers a wealth of information to predict functional interactions, including data from the literature that are used to curate a resource like Complex Portal, our model exclusively exploited co-evolutionary and co-expression features. Indeed, when considering only coevolution and co-expression scoring from String, our model achieved better performances (Figure 3C). On the IA direct interaction set, our model outperformed all the different scoring criteria from String (Figure 3D).

We inspected the importance of the features in the different models. Consistently with the feature exploratory analysis (see above), we found that in the task of stable complexes interaction classification the most important features are dominated by co-expression, followed by co-evolutionary features (Figure 3E). On the other hand, in the tasks of direct interaction prediction, the most important features are the co-evolutions of interacting domains or entire genes (Figure 3F). This suggests that while for the prediction of stable complexes the cellular contextual information, expressed as gene co-expression, is critical to achieve good prediction results, for the prediction of direct interactions molecular and structural features, such as co-evolution of interacting domains, are more important.

### Human proteome-wide prediction and discovery of novel complexes

We employed the CP models to evaluate their capability to recover known complexes from Complex Portal. In this respect, we predicted with the models trained on Complex Portal the probability of interaction of every possible protein-protein pair within the human proteome. We then used the probabilities returned by the models as weights to obtain a proteome-wide, weighted adjacency matrix. We clustered the resulting graph using the Louvain approach (Nguyen et al., 2008) to retrieve communities of interacting proteins. We then assessed the recovery of known complexes from Complex Portal within detected modules at different clustering depths using the geometric accuracy, a cluster comparison metric specifically developed for protein-protein interaction networks (Brohée & van Helden, 2006) (Figure 4; see Methods). We found that our approach outperformed all String baselines, including String and String Physical networks, in recovering known complexes on the entire CP sets (Figure 4).

**Figure 4.**
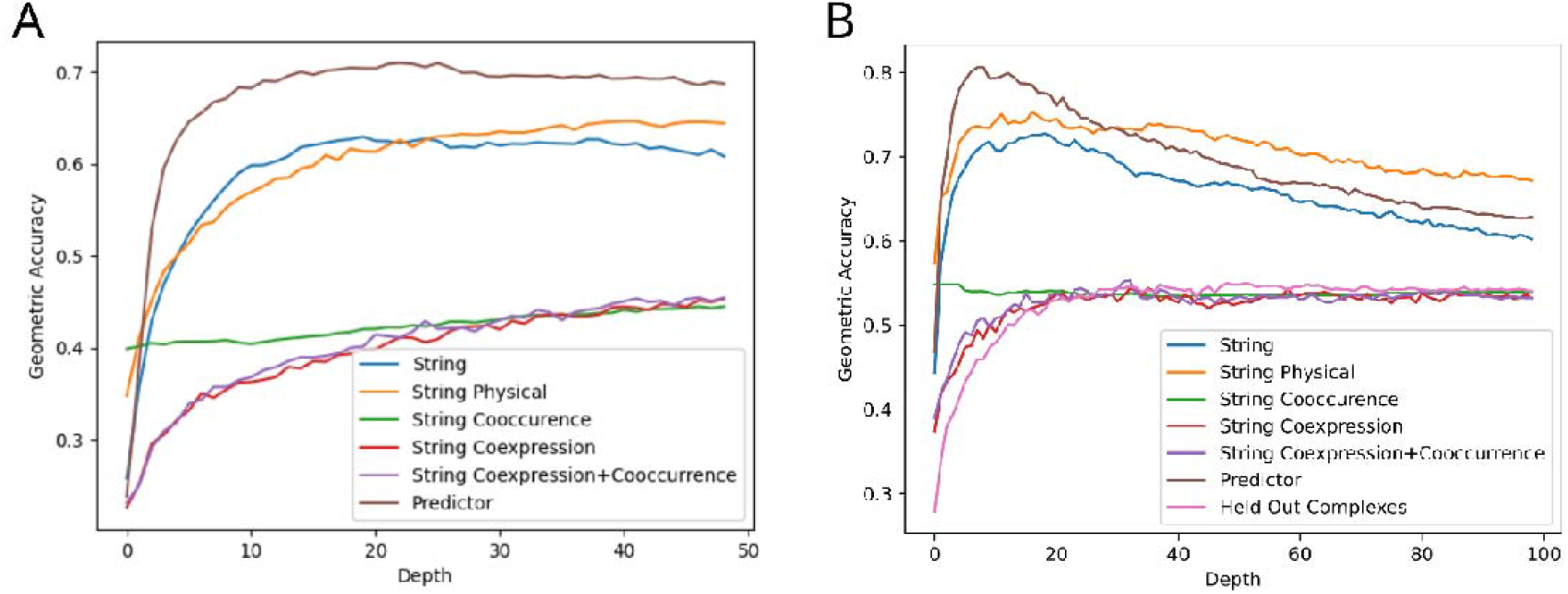
Complex recovery at a proteome scale using the STRING and the predicted networks. Complex recovery is assessed using Louvain clustering at different depths: A) Performance evaluation on the network of proteins belonging to at least one known complex; B) Performance evaluation on the network of proteins belonging to the complexes that were held out from the training set of the stable complex prediction model in pink.

## Discussion

In this study we have developed machine learning models to accurately predict interactions either involved in direct associations or mediating the formation of stable complexes. We demonstrated that by leveraging co-evolution information, such as the co-presence in sequenced genomes of gene or interacting domain pairs, as well as the information of gene co-expression from both healthy and cancer tissues, we could achieve predictive performances competitive with state-of-the-art methods such as String (Szklarczyk et al., 2023). Notably, the String scores obtained with similar features to the ones we employed, i.e. co-presence and co-expression scores, performed much worse.

Intriguingly, co-expression information, particularly from cancer tissues, is more important for the prediction of stable complexes, suggesting that contextual information is important for the definition of stable complexes. On the other hand, we found that co-evolutionary information, particularly the one of domain-domain pairs known to form 3D interfaces, was deemed more important for the prediction of direct interactions, which are expected to involved 3D structured interfaces.

Both of our models showed better performance than String in recovering known complexes when clustering the adjacency matrix of the human proteome obtained by weighting the edges via the interaction probabilities obtained from the models.

Taken together, these results suggest that our model could be used to process any given input interactomic dataset to discriminate interactions more likely to be either direct or involved in stable complexes, from other interaction types. The model can also potentially be applied to the prediction of protein complex topologies as well as to the discovery of new complexes at a proteome-scale level. This method could be pipelined with structural prediction algorithms such as AF-multimer to narrow down the list of candidates to model as well as to suggest more likely higher order complex topologies.

## Methods

### IA Dataset

The IA set was derived from the IntAct database (May 2024 release; https://www.ebi.ac.uk/intact/download/ftp), considering only interactions involving human proteins. Interactions were classified into four categories based on IntAct’s “interaction type”:

- “association”: included interactions labeled as “colocalization,” “proximity,” or “association.”
- “physical association”: retained the original “physical association” label.
- “non-enzymatic direct interaction”: included interactions labeled as “direct interaction.”
- “enzymatic direct interaction”: included interactions annotated with the name of an enzymatic reaction.

In cases where multiple instances of the same protein pair existed in IntAct, these were aggregated, and the interaction was assigned the highest priority class from the following order: “enzymatic direct interaction” > “non-enzymatic direct interaction” > “physical association” > “association”. In the end the dataset contained 421,579 PPI: 302,139 associations, 111,368 physical associations, 5,894 non-enzymatic direct interactions and 2,178 direct interactions.

We also annotated as Complex Portal interactions, the protein pairs that were included in the same complex in at least one of the instances of Complex Portal (9,654 positives).

### CP Dataset

The CP dataset was generated taking as positive instances all protein pairs that co-occur within the same Complex Portal(Meldal et al., 2022) complex (15,881 pairs). Negative instances were randomly sampled from the human proteome, excluding positive pairs, to a total of 674,637, to reproduce the same positive/negative ratio of the IA dataset.

### Coevolution based features

For each protein of the human proteome, we extracted the list of genomes containing an orthologous sequence of that protein from the Orthologous MAtrix (OMA) database (Altenhoff et al., 2021) using a custom Python script using the PyOMADB client (Kaleb et al., 2019) (December 13, 2023, database version July 2023).

Coevolution between each protein pair was measured using two approaches:

- The Jaccard similarity coefficient between the sets of genomes containing orthologs of the two proteins.
- The mutual information regarding the presence/absence of the two proteins across genomes.

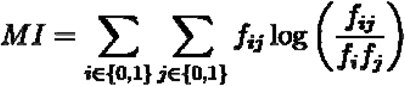

Where, i = 1 (or 0) denotes the presence (or absence) of an ortholog of the first protein, and j = 1 (or 0) denotes the presence (or absence) of an ortholog of the second protein. ‘fij’ represents the frequency of observing the combined state (i, j).

### Domain coevolution

We retrieved domain annotations for each protein from the InterPro database (Blum et al., 2021) (v101.0). For a given domain D within a protein P, we define the set of genomes containing an ortholog of D as those genomes where an ortholog of P exists and also possesses a domain with the same ID as D. This definition allows us to measure the coevolution between two domains in a manner analogous to protein coevolution:

- Using the Jaccard similarity coefficient between the sets of genomes containing orthologs of the two domains.
- Calculating the mutual information regarding the presence/absence of orthologs of the two domains across genomes.

For our classifier, we utilize the mean coevolution of all domain pairs as features. This means that coevolution is computed using both the Jaccard similarity coefficient and mutual information approaches.

### Interacting domains

We compiled a catalog of domain pairs with at least one structurally resolved interaction interface in the PDB. We adopted InterPro’s domain definitions and considered two domains to be in structural contact if they had a minimum of 5 residue-residue contacts. A residue-residue contact was defined as having a distance of 8 Å or less between the Cβ atoms (or Cα for glycine) of the two amino acids.

For our classifier, we used as additional features the mean coevolution of the domain pairs that have at least one structurally resolved interchain interaction interface between domains with the same IDs. In cases where no such domain pairs existed, these features were set to 0.

### Co-expression based features

Bulk RNA-seq data is obtained from UCSC Xena(Goldman et al., 2020). The cohort includes data from 3 projects: TCGA, focusing on cancer tissue; GTEx, healthy tissue and TARGET, pediatric data. All data from TARGET is removed from subsequent analysis, and only primary tumor data is kept from TCGA. The resulting dataset consists of a total of 18305 samples, 7775 healthy tissue samples from GTEx and 10530 primary tumor samples from TCGA.

First, outliers are identified and removed based on broad quality control metrics. Samples with less than 20000 or more than 40000 genes detected are removed, same for samples with less than 10 millions or more than 120 millions total counts. Samples in which the top 100 most expressed genes accounted for more than 90% of total counts were also removed.

The data is then split according to tissue and histology (e.g. healthy pancreas and pancreatic adenocarcinoma are considered different tissues) and tissue-specific preprocessing is applied. In particular, we consider the following quality control metrics: logarithm of total counts, logarithm of genes with at least one count (in both cases a pseudocount is added to handle 0 values) and percentage of counts in the top 100 most expressed genes. Only samples in which all QC metrics are within 5 median absolute deviations from their median value of the tissue are kept.

Next, we remove lowly expressed genes to reduce the effects of sparsity on the correlation analysis and prevent the formation of clusters of lowly expressed genes, which could complicate interpretation of the results. To this end, genes with less than 15 counts in more than 75% of the samples are removed from each tissue.

We then apply normalization and variance stabilizing transformation from DeSeq2 (Love et al., 2014) (PyDeSeq2 implementation (Muzellec et al., 2023)). Finally, we use principal component analysis to identify additional outliers. Specifically, we compute the distance of each point from the centroid in the principal components space (considering the first 10 principal components). Samples that are more than 5 standard deviations away from the centroid are then removed.

In order to generate co-expression based features, we use the implementation of the WGCNA (Langfelder & Horvath, 2008)algorithm provided in the PyWGCNA Python module (Rezaie et al., 2023).

WGCNA operates by first computing the Pearson correlation coefficient of each pair of genes across all samples, the correlation is then processed to obtain a non-negative, weighted, undirected adjacency matrix (signed adjacency):

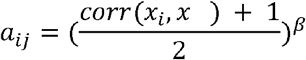

Where *β* represents a soft thresholding power and its purpose is thresholding the adjacency without actually binarizing it. The optimal value of *β* is determined according to the approximate scale-free criterion (Zhang & Horvath, 2005) selecting a target value of R^2^ = 0.85.

Finally, the TOM matrix is obtained from the adjacency matrix (Zhang & Horvath, 2005):

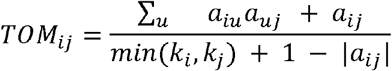

Where k_*i*_ = Σ_*j*_ *a*_*ij*_ is the degree of gene i in the co-expression network.

### Model training

To classify the protein pairs in the different interaction types we trained an XGBoost classifier using the official package (Chen & Guestrin, 2016). Hyperparameters optimization was performed with Optuna (100 trials), maximizing average precision. All 164 previously described features were used as input. Undefined features (that were impossible to compute for certain pairs) were set to −2 (a value outside of the range of all the features), and pairs with all features undefined were removed. Both the IA and CP datasets were split into 80% training, 10% validation (used for hyperparameters optimization), and 10% testing sets. The final models were trained on the combined training and validation sets before evaluation on the held-out test set.

The stability of the models was evaluated with a 10-fold cross-validation using the scikit-learn python package (Pedregosa et al., 2011).

### Testing on held out complexes

We randomly selected 10% of the original complexes as Test Complexes and another 10% as Validation Complexes. The training set included all pairs in the CP set that did not involve proteins from either the Test or Validation Complexes. The validation set, used for hyperparameter optimization, included all pairs in the CP set exclusively involving proteins found in the Validation Complexes but not in the Test Complexes. After hyperparameter optimization, the model was trained on all pairs in the CP set that did not involve proteins from the Test Complexes. It was then tested on all possible pairs of proteins found within the union of all the Test Complexes (including those not present in the CP set).

### STRING scores

We benchmarked our models’ predictions against evidence scores from STRING (v12.0), mapping UniProt protein IDs to STRING IDs using the UniProt ID mapping files (Huang et al., 2011). We considered both the ‘combined score’ and the physical networks from STRING. Additionally, we included cooccurrence and coexpression scores for comparison, as these reflect the types of features utilized by our predictor.

### Clustering

To identify clusters of proteins likely to form complexes, we constructed a network of all proteins belonging to at least one complex in the Complex Portal. We used the predicted probability of two proteins belonging to the same complex as the edge weight between them in this network. Clusters were then identified within this weighted network using the Louvain algorithm, using resolution ranging from 1 to 100. Clustering was performed using NetworkX (v3.2.1; https://networkx.org/) and CDlib library (v0.4.0; https://cdlib.readthedocs.io/en/latest/). The clusters obtained at each resolution were compared to the known complexes using the cdlib.evaluation.geometric_accuracy function.

## Software

Plots were generated with customized scripts in python (v3.9.13), using matplotlib (v3.5.1) and seaborn (v0.11.2). Statistical analysis was performed using scikit-learn (v1.0.2).

## Acknowledgement

We gratefully acknowledge computational resources of the Center for High Performance Computing (CHPC) at SNS.

## Notes

### Competing Interest Statement

The authors have declared no competing interest.

